# The “two-brain” approach reveals the active role of task-deactivated default mode network in speech comprehension

**DOI:** 10.1101/2021.03.02.433669

**Authors:** Lanfang Liu, Hehui Li, Zhiting Ren, Qi Zhou, Yuxuan Zhang, Chunming Lu, Jiang Qiu, Hong Chen, Guosheng Ding

**Author notes:** **Corresponding author:** either Guosheng Ding, State Key Laboratory of Cognitive Neuroscience and Learning, Beijing Normal University, No. 19 Xinjiekouwai Street, Beijing 100875, China, or Hong Chen, School of psychology, SouthWest University, No. 2 Tiansheng Street, Beibei, Chongqing, 400715.

## Abstract

During speech comprehension, as listeners need to keep tracking the external audio streams, the default mode network (DMN) is often de-activated and anticorrelated with task-positive networks. Such a pattern has been interpreted as the suppression of the DMN to support externally-oriented cognitive processes. Taking a “two-brain” approach, the current study demonstrated that, despite exhibiting deactivation and anticorrelated with the language network and executive control network, the DMN was not suppressed but played an active role in spoken narrative comprehension. This was evidenced by significant listener-speaker neural couplings in both the posterior and anterior DMN and the positive correlation between the coupling strength and listener’s speech comprehension. Moreover, we demonstrated that the functionality of posterior DMN depended on its interaction with the executive control network, rather than its level of activation. Finally, Dynamic Causal Modeling together with the two-brain results indicates the language and executive control networks, the anterior DMN, and the posterior DMN occupied the bottom, intermediate and top layers of a hierarchical system, respectively. These results suggest that the DMN may primarily serve as an internally-oriented system that cooperates with the externally-oriented networks, which may allow the transformation of external acoustic signals into internal mental representations during language comprehension.

## 1. Introduction

In neurolinguistic studies, verbal communication has been usually approached as a signal transmission process, and the main attention has been paid to the neural responses bound to and triggered by the occurrence of speech signals (Stolk et al. 2016). Within this signal-centered framework, the crucial role of a set of inferior frontal and temporal regions (i.e., the language network, LN) in coding and decoding the sensory-motor, syntactic and semantic properties of speech signals, as well as a set of middle frontal and parietal regions in monitoring and coordinating these processes (i.e., the executive control network, ECN) (Ye and Zhou 2009), have been well recognized. Nevertheless, in real-life situations, the speech signals we receive are usually impoverished or ambiguous. Yet, we can still decipher the meaning intended by a speaker accurately. By contrast, current state-of-art artificial agents, such as Apple’s Siri or Microsoft’s Cortana, often make communicative errors despite they have mastered the signal decoding rules. These observations indicate that successful speech understanding demands more than signal decoding.

Indeed, psychological models have long proposed that, in addition to the perceptual and linguistic analyses, to successfully understand the speaker’s mind during verbal communication engages a set of extra-linguistic processes, mainly including mentalizing, self-projection and the theory of mind (Carruthers and Smith 1996; Zwaan and Radvansky 1998; Frith and Frith 1999). These processes have been frequently attributed to the default mode network (DMN) (Spreng et al. 2008; Spreng and Grady 2009; Li et al. 2014), an intrinsically organized functional system composed of the posterior cingulate cortex, precuneus, medial prefrontal cortex (mPFC), and lateral parietal cortex.

Nevertheless, current empirical evidence regarding the functional role of DMN in speech processing seems conflicting. On the one hand, as auditory language processing requires participants to keep tracking external audio streams, the DMN has been repeatedly found to be deactivated relative to low-level baselines (Wilson et al., 2007; Szaflarski et al., 2012; Rodriguez Moreno et al., 2014; Horowitz-Kraus et al., 2017; Cuevas et al., 2019), and anticorrelated with networks dedicated to externally-oriented processes (including the LN and ECN) (Fox et al. 2005; Uddin et al. 2009; Smith et al. 2012). The task-induced deactivation and network anticorrelation seem to imply that the DMN was “suppressed” to support externally-oriented cognitive activities (Anticevic 2012; Gauffin et al. 2013; Zhou et al. 2016).

On the other hand, studies measuring the consistency of blood-oxygen-level-dependent (BOLD) signal fluctuations across subjects experiencing the same narratives (i.e., intersubject correlation, ISC) have found significant ISC in the DMN regions (Wilson et al., 2007; Lerner et al., 2011; Simony et al., 2016; Yeshurun et al., 2017). In addition, the ISC in the DMN was modulated by the manipulation of story belief (Yeshurun et al., 2017) and the coherence of the stimuli’s temporal structure over minutes of time (Lerner et al., 2011; Simony et al., 2016). These results are postulated to reflect the active role of DMN in semantic/conceptual processing.

How to reconcile a potentially active role of the DMN as argued by ISC studies with the longstanding observation that the DMN shows task-induced deactivation and anticorrelated with task-positive networks? One possible explanation is that the significant ISC (and its change) in the DMN reported in previous studies is an epiphenomenon of task disengagement, rather than signifying a positive role. It is well known that activities in DMN regions are systematically modulated by task difficulty (Zwaan and Radvansky 1998; McKiernan et al. 2003; Greicius and Menon 2004; Humphreys et al. 2015). If subjects find the same part of the narrative more or less engaging, the DMN will be down- or up-regulated similarly across subjects, which would result in the correlation of brain signal fluctuations across subjects (Wilson et al. 2007). Alternatively, the DMN is indeed critically engaged in language comprehension, but the processes (likely internally-directed) it subserves are opposite to those (likely externally-directed) subserved by the task-positive networks. This hypothesis is put forward mainly based on the linguistic theory that the basic design for language faculty includes not only externally-directed modules for sensory-motor and linguistic processing but also an internally-directed module for conceptual-intentional representations (Berwick et al. 2013). However, it remains to be established that the DMN showing deactivation and anticorrelation with task-positive networks is not “suppressed” but actively involved in language comprehension. Moreover, how the task-activated and task-deactivated networks are organized to support speech comprehension remains largely unknown.

To address the above issues, we took a “two-brain” approach and combined within-brain and between-brain analyses. In our experiment, a speaker telling real-life stories and later a group of listeners hearing the stories were scanned with fMRI. Rather than relying on the ISC across listeners to determine the involvement of DMN, we focused on neural couplings (i.e., the correlation of brain activities over time) between the listener and the speaker. Compared to the ISC, the inter-brain coupling may have at least two advantages to uncover the functional role of DMN in task processing. First, the cognitive meaning of listener-speaker neural coupling might be clearer than the ISC. Accumulating evidence suggests that inter-brain couplings at different cortical regions reflect the alignment between communicators at different levels of mental representations (Menenti et al. 2012; Schoot et al. 2016; Stolk *et al.* 2016). Indeed, it has been found that stronger inter-brain coupling was associated with better communication (Stephens et al. 2010; Liu et al. 2020; Li et al. 2021; Liu et al. 2021). Second, the neural coupling between a speaker-listener pair is likely less susceptible to the influences of task difficulty, since speech production is typically more cognitively demanding than speech comprehension (Hendriks 2014).

The group Independent Component Analysis (ICA) was applied to detect large-scale neural networks and the associated network time series. Compared to selecting regions of interest, this data-driven approach has been shown to render better representations of neural networks and produce more reliable results (Guo et al. 2012; Yu et al. 2017). As accumulating evidence showed that the DMN is not a homogenous network, but generally composed of two functionally heterogeneous subsystems (Uddin *et al.* 2009; Seghier and Price 2012b), the anterior and posterior parts, we examined the anterior DMN (aDMN) and posterior DMN (pDMN) separately. The two externally-oriented networks, the LN and ECN, were also investigated.

We first examined the activation pattern and network functional connectivity within the listeners’ brains to compare with previous “single-brain” studies. Next, we examined listener-speaker neural coupling and its association with task performance. Upon establishing the active role of DMNs in speech comprehension, we explored the organization of network communication applying Dynamic Causal Modelling (DCM) to gain insight into the mechanism of the DMN’s contribution to language processing. Finally, we tested whether the functionality (measured by the inter-brain coupling) of DMN depends on its interaction with other networks or on its response amplitude to stimuli.

## 2. Materials and Methods

### 2.1 Participants and experimental procedure

A total of 69 Chinese college students (aged 19-27 years) participated in this study, including one female speaker and 68 listeners (35 females). We also recruited a native Mongolian speaker (a female college student) to serve as a control condition. All participants were right-handed and reported no history of physiological or mental disorder. None of these listeners had learned Mongolian. The datasets of four participants listening to the Chinese story and four participants listening to the Mongolian story were excluded from further analysis due to excessive head movements during the scanning (more than 3mm or 3 degrees). Another two participants were excluded due to their extremely low comprehension scores (lower than 3.5 standard deviation from the group mean), which indicated they probably did not listen to the story attentively. Written informed consent was obtained from all participants under a protocol approved by the Reviewer Board of Southwest University in China.

We first collected the data from the two speakers telling stories based on their personal experience (10 min for each) when undergoing fMRI scanning. A noise-canceling microphone (FOMRI-III, Optoacoustics Ltd., Or-Yehuda, Israel) was used to simultaneously record the speech. After further offline denoising by Adobe Audition 3.0 (Adobe Systems Inc., USA), the audio recordings were played back to each listener during scanning. To make the listeners attend to the stories, we informed them beforehand that a test about the content of the story would be given after the scanning. During the scanning, the same set of visual stimuli were presented to the speakers and listeners, consisting of a fixation cross lasting for 20s and then an icon of a horn lasting until the end of the scanning run (Fig. 1A). The horn was designed to prompt the participant to start speaking or listening immediately. To control for the effect of low-level acoustic processing, we also played the story told by the Mongolian speaker to each listener.

**Figure 1.**
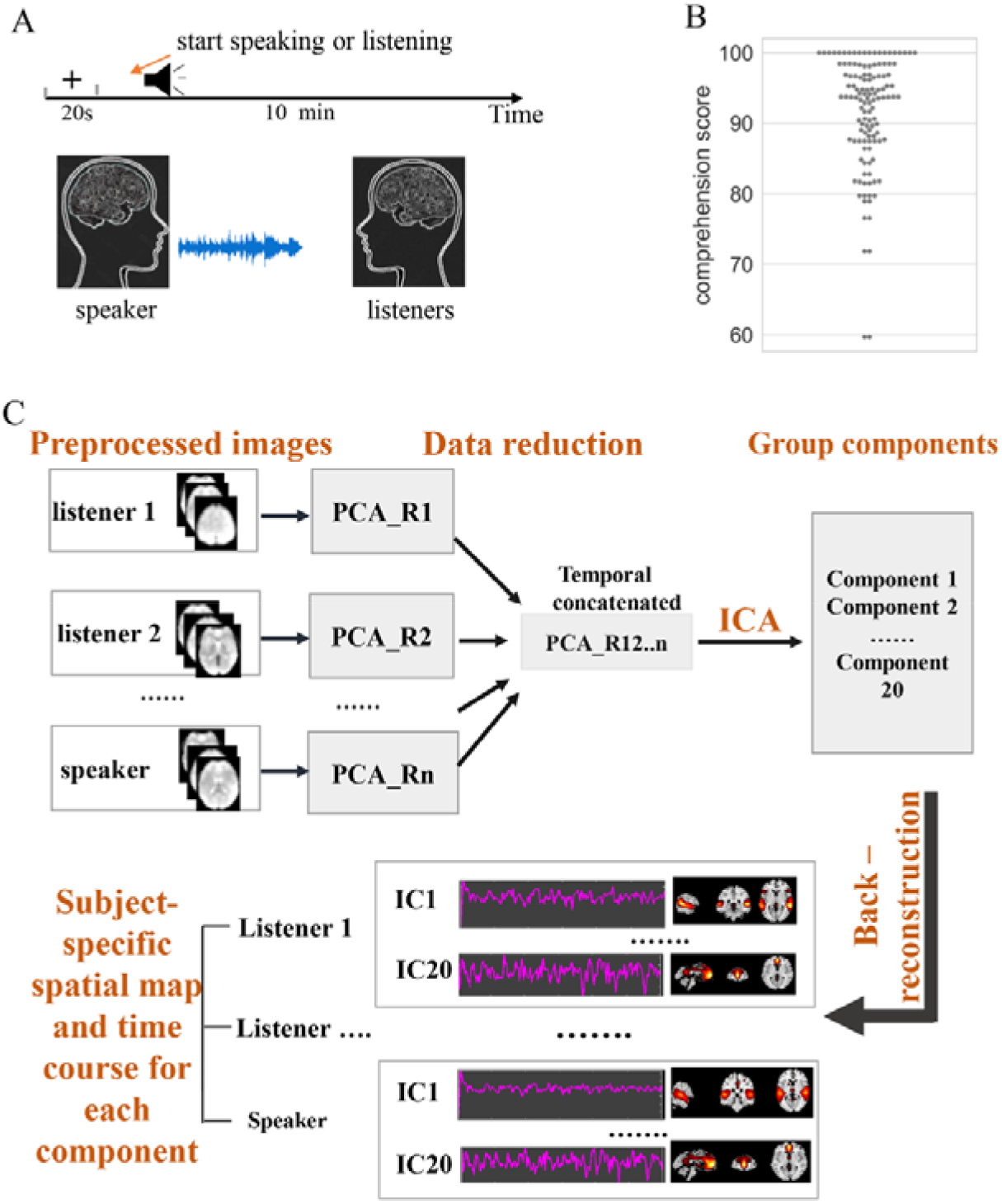
Experimental design, behavioral performance and data analysis. (A) One speaker told real-life stories while undergoing fMRI scanning. The audio recordings of the stories were then played to a group of listeners (N=62) during fMRI scanning. (B) The distribution of comprehension scores for the story across the listeners. (C) The procedure of Group Independent Analysis.

### 2.2 Behavioral assessment

To assess the degree to which the listeners understood the speech, we conducted an interview with each listener at the end of the scan. In the interview, the listeners were required first to retell the story with as much detail as possible. Next, experimenters asked the listeners several questions concerning the part of the contents not present in their free recalls. Two independent raters then scored the listeners based on the audio recordings of the interview. Details about the scoring procedure were provided in the supplementary material. The two raters had a high agreement on their assessments (r_(64)_ = 0.80, by Pearson’s correlation), thus their assessments were averaged to be used as the measurement for speech comprehension. We also measured participants’ memory spans using a digit memory test (Wechsler 1987) to account for a potential effect of memory capacity on the story-retelling task. The number of digits correctly repeated forward and backward were used as two covariates in the following brain-behavior analysis.

### 2.3 MRI acquisition and preprocessing

Imaging data were collected with a 3T Siemens Trio scanner in the MRI Center of the Southwest University of China. Functional images were acquired using a gradient echo-planar imaging sequence with the following parameters: repetition time = 2000 ms, echo time = 30 ms, flip angle = 90°, field of view = 220 mm^2^, matrix size = 64 × 64, 32 interleaved slice, voxel size = 3.44 × 3.44 × 3.99 mm^3^. T1 structural images were acquired using a MPRAGE sequence with the following parameters: repetition time = 2530 ms, echo time = 3.39 ms, flip angle = 7°, FOV = 256 mm^2^, scan order = interleaved, matrix size = 256 × 256, and voxel size = 1.0 × 1.0 × 1.33 mm^3^.

After discarding the first three volumes to allow for the equilibration of magnetic fields, a total of 310 volumes were collected in each run. The DPABI toolkit (Yan et al. 2016) based on SPM12 (www.fil.ion.ucl.ac.uk/spm/) was utilized for image preprocessing. The preprocessing steps included slice-timing correction, spatial realignment, co-registration to individual subject’s anatomical images, normalization to the Montreal Neurological Institute (MNI) space, resampling into a 3 × 3 × 3 mm^3^ voxel size, and smoothing (FWHM = 7mm). The resultant images were further detrended, nuisance variable regressed, and high-pass filtered (1/128 Hz). The nuisance regression included the removal of five principal components of white matter and cerebrospinal fluid within individual subjects’ T1 segmentation mask (Behzadi et al. 2007), as well as Friston’s 24 motion parameters (including each of the six motion parameters of the current and preceding volume, plus each of these values squared) (Friston et al. 1996).

### 2.4 Data analysis

#### 2.4.1 Group Independent Component Analysis and component selection

For the successful and failed communication conditions separately, a group spatial ICA was conducted using the Group ICA of fMRI Toolbox (GIFT). Before entering the analysis, the first 10 volumes of the preprocessed functional image corresponding to the 20s resting (fixation) period were discarded to avoid the potential modulation of state change on the spatial patterns of components (Calhoun et al. 2008). The dimensionality of preprocessed images from each participant was first reduced to 30 dimensions using principal component analysis. Next, the dimension-reduced data from each participant (including both the speaker and all listeners) were temporally concatenated and a group dimension reduction was performed. Then, twenty independent sources were generated from the group data with the ICA using the infomax algorithm. We choose to separate the whole brain signals into 20 independent components because this dimension can well detect those typical brain networks meanwhile keeping high stability of ICA estimation (Pamilo et al. 2012). Finally, spatial maps and associated time series were reconstructed for each participant based on the aggregate components and original data via spatial-temporal regression (Beckmann et al. 2009) (Fig. 1C). This dual-regression technique has been demonstrated to show high test-retest reliability (Zuo et al. 2010) and is widely used to explore the functional connectivity of brain networks. Since the spatial maps and time series of components have arbitrary units after the back-reconstruction step, they were scaled to z-scores. Those time series associated with network components were used in the following within-brain and between-brain connectivity analyses. A one-sample *t*-test was conducted on individual subjects’ component maps to obtain the representative spatial distribution of each component. To test the robustness of our findings, we also separated the whole brain signals into 30 independent components and re-performed the analyses (supplementary material).

The four networks of interest, as well as the control network, were identified based on the spatial overlap of the component map (i.e., the *t*-statistic map) with the meta-analytic maps generated by the Neurosynth (https://neurosynth.org). The spatial overlap of the two types of maps was quantified by the Dice score (Dice 1945). Details for the calculation of the Dice score were provided in the supplementary material. The term “default mode network” was adopted to search for the meta-analytic map of the DMN. The terms “language” and “executive control” were adopted to search for the meta-analytic maps of LN and the ECN, respectively. The meta-analytic map of the visual network was identified by the search term “visual cortex”. Upon identifying those components corresponding to the five networks, we thresholded the *t*-statistic maps with a *p* < 0.05 (corrected for multiple comparisons with family-wise error (FWE) correction) for visualization and making masks used in the bellowing activation analyses.

#### 2.4.2 BOLD signal change

To assess the modulation of network activity by external stimulation, we measured percent changes in BOLD signal amplitudes during speech listening (the task phase) relative to the 20s fixation period (the baseline phase). The task onset was shifted forward by 6s to compensate for delays in the brain’s hemodynamic response. The BOLD signal changes were defined as [(*task-baseline*) / *baseline*], where *task* and *baseline* denote the mean of time points within the task phase and the baseline phase, respectively. We first computed the voxel-wise signal change for each subject. Then for each network, the mean value across voxels within the thresholded component map was calculated and a *t*-test was applied to examine whether the mean signal change in the network differed from zero. In addition to using the time courses of single voxels, we also estimated the BOLD signal changes based on representative network time courses derived from a separately-conducted group ICA. Details about this analysis were provided in the supplementary material.

#### 2.4.3. Within-brain function connectivities

For each listener, functional connectivities between networks were assessed by Pearson’s correlation using the component time courses derived from the ICA. The r values were converted to Fisher’s z values and analyzed using *t*-tests.

#### 2.4.4 Measurement for listener-speaker neural coupling

The listener-speaker neural couplings were measured by Pearson’s correlation coefficients between the ICA-derived network time courses from each listener’s brain and the time courses of the homologous network from the speaker’s brain. The r-values were transformed to z-values using Fisher’s z transformation to normalize their distribution. Previous work has demonstrated that listeners need time to process the high-level information in speeches in order to get aligned with the speaker (Stephens *et al.* 2010; Kuhlen et al. 2012; Liu et al. 2017; Liu *et al.* 2020). To account for this effect, we repeated the inter-brain correlation analysis by shifting the time courses of listener’s network activity relative to that of the speaker by 0-14s with 2s increments. At shift zero, the listener’s network activity was time-locked to the speaker’s vocalization. Two-tailed *t*-tests were applied to assess whether the neural couplings differed significantly from zero at the group level. Multiple comparisons were corrected with an FDR q = 0.01 using the Benjamini–Hochberg procedure (Benjamini and Hochberg 1995).

#### 2.4.5 Brain-behavioral correlation

The Pearson’s partial correlation analysis was applied to assess the correlation between the coupling strength of each network and listeners’ comprehension scores, controlling for the potential effect of memory span. To evaluate the generalizability of the brain-behavior relationship we may find, we further performed within-data set cross-validation (Shen et al. 2017). Specifically, a linear regression model was built on the data from all participants but one, taking inter-brain couplings in the networks which showed significant brain-behavior correlations as the predictors and listener’s comprehension score as the outcome. Then the model was used to predict the comprehension score of the left-out participant based on his or her neural coupling with the speaker. The significance of the prediction was assessed via permutations. In each permutation (total N = 1000), we randomized the comprehension scores across the training sample and re-conducted the prediction. A *p*-value was calculated as the percentage of permutations that showed a lower r-value than the actual value.

#### 2.4.6 Dynamic Causal Modeling for network communication

Functional connectivities only capture the statistical dependencies between two signals (Friston 2011). To gain insights into the organization of functional communication among the four networks during speech comprehension, we further conducted Dynamic Causal Modeling analysis using the DCM toolbox in SPM12. Given previous reports that language is processed in the brain in a hierarchal manner (Davis and Johnsrude 2003; Lerner et al. 2011; Ding et al. 2016), we presumed that the four networks were organized into layers. Four models consisted of three layers with varying components in those layers were tested. Model 1 hypothesized the two externally-directed networks (the LN and ECN) lied in the same layer, having causal interactions with the aDMN in the intermediate layer, which in turn having causal interactions with the pDMN. Model 2 differed from model 1 in that the positions of aDMN and pDMN were exchanged. Model 3 hypothesized the two DMN subsystems lied in the same layer, having causal interactions with the ECN in the intermediate layer, which in turn having causal interaction with the LN. Model 4 varied from model 3 in that the positions of ECN and LN were exchanged. All models included recurrent connections of each network and had no input entering the system. All connections in these models were bidirectional. Since the speech signals presented during the task were continuous stimuli, we fitted models based on the frequency-domain cross-spectral density of the time courses, which is a technique originally developed for modeling resting-state fMRI data (Friston et al. 2014). The model with the highest probability was identified through random-effect Bayesian Model Selection.

#### 2.4.7 Analysis for the association between within-brain network BOLD signal change and functional connectivity and listener-speaker neural coupling

Finally, we explored whether the functionality of DMN depends on its interaction with other networks or on its response amplitude to stimuli. For the pDMN and aDMN separately, we calculated the Pearson’s correlation between inter-brain neural coupling and network mean signal changes and functional connectivity within listeners’ brains.

## 3. Results

### 3.1 Behavioral results

On the story-recall task administered shortly after the scanning, participants scored on average 91.58 ± 8.00% (Fig. 1B). For the Mongolian story (the control condition), the interview showed that the participants had tried to comprehend the speech but all failed.

### 3.2 ICA-based network detection

By measuring the spatial overlap of each ICA component with the meta-analytic maps derived from the Neurosynth, we identified four networks of interest and a visual network serving as a control (Fig. 2A). Two components consistently showed high Dice scores with the meta-analytic map of DMN. The first component was mainly comprised of the bilateral medial frontal gyrus, insular, precuneus, and angular gyrus, peaking at the ventral mPFC (MNI coordinates: 0, 57, 3). This component matches with the aDMN in the literature (Xu et al. 2016). The second component covered mainly the bilateral precuneus, inferior parietal lobe, and part of the medial frontal gyrus, peaking at the left precuneus (MNI coordinates: −3, −72, 33), which matches the pDMN in the literature (Xu *et al.* 2016). The LN component covered mainly the bilateral superior and middle temporal, precentral, and inferior frontal gyrus, as well as the supplementary motor area, peaking at the right superior temporal gyrus (MNI coordinate: 57, −24, −3). The ECN component was primarily comprised of the bilateral middle/inferior frontal gyrus and the superior/inferior parietal lobule, peaking at the right inferior frontal operculum (MNI coordinates: 51,15, 27). The control (visual) network covered mainly the bilateral inferior and middle occipital gyrus, fusiform and lingual gyrus, peaking at the left middle occipital gyrus (MNI coordinate: −24, −96, 0). The Dice scores and the spatial maps of other components derived from the group ICA were presented in the supplementary material (Fig. S1 and Table S1).

**Figure 2.**
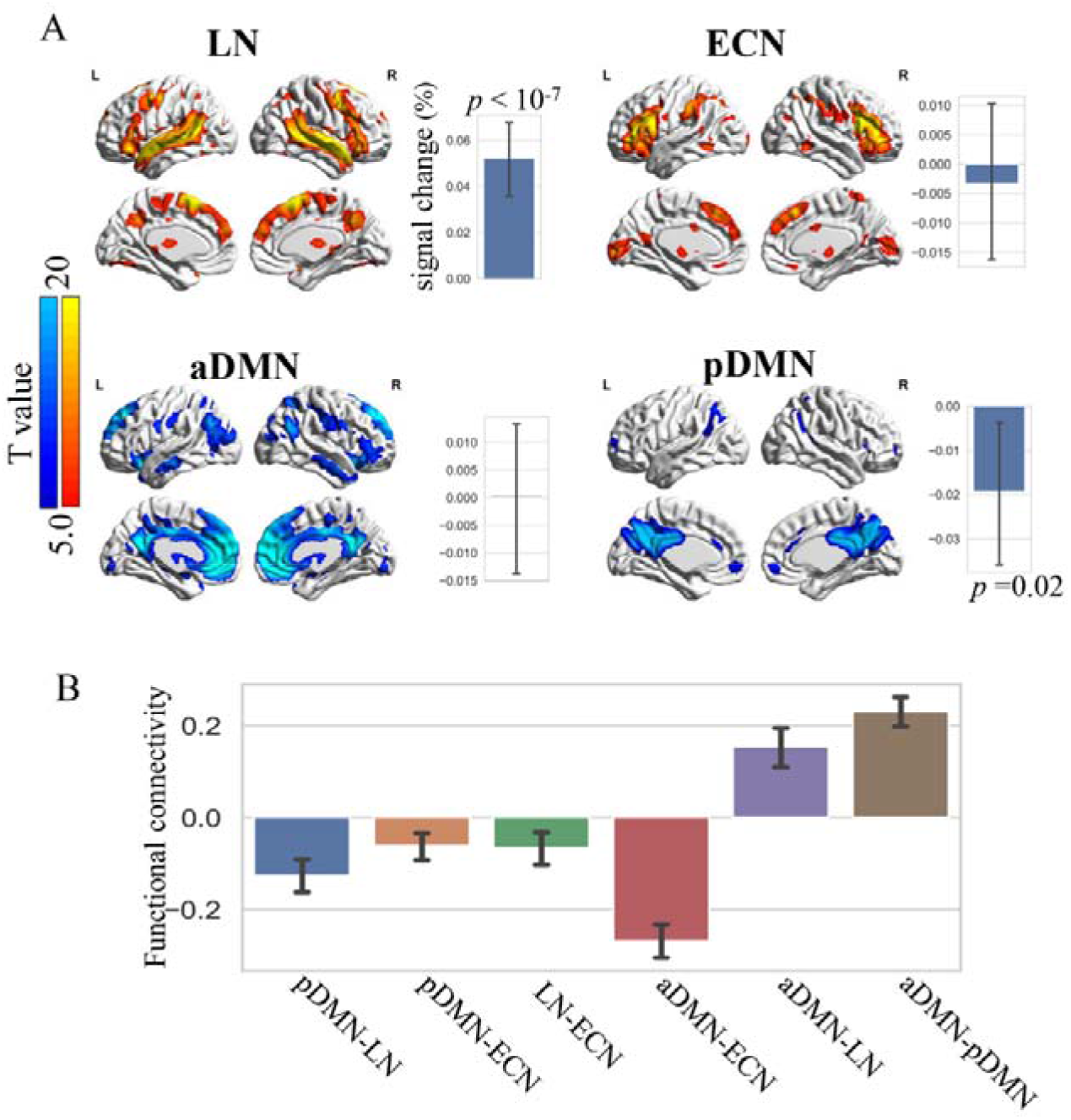
Network activation and functional connectivity pattern. (A) The spatial distribution of networks derived from the group ICA and their activation patterns. (B). Functional connectivities between networks. All connectivities were significant with an FDR-corrected *p* < 0.01. The error bar denotes the 95% confidence interval of the group mean. LN: language network; ECN: executive control network; aDMN: anterior default mode network; pDMN: posterior default mode network.

#### 3.3 Within-brain activation pattern

In line with the previous studies (Seghier and Price 2012a; Rodriguez Moreno et al. 2014), *t*-tests on the mean signal change over voxels in each of the four networks revealed significant activations in the LN (*t*_(61)_= 6.21, *p* < 10^−7^) and significant deactivation in the pDMN (*t*_(61)_ = −2.35, *p* = 0.02) during speech listening relative to the fixation period. No significant signal change was observed in the aDMN (*t*_(61)_ = −0.05, one-tailed *p* = 0.96) or the ECN (t_(61)_ =-0.46, *p* = 0.64) (Fig. 2A). Consistent results were obtained when the signal change was assessed using the time courses derived from the group ICA (Fig. S2).

We note the baseline used to calculate the BOLD signal change might be too short. To assess this effect, we compared this method with a conventional General Linear Modeling (GLM) method which included longer baselines using an independent dataset. The results showed that the activation and deactivation patterns obtained by the two approaches were similar, with the one obtained from the GLM being statistically more significant (Fig. S3). Thus, a relatively short baseline does not compromise the findings of opposite activation patterns between the pDMN and LN.

#### 3.4 Within-brain functional connectivity

Consistent with previous studies (Uddin *et al.* 2009; Smith *et al.* 2012), we observed negative functional connectivities of the pDMN with both the LN (t_(61)_=-6.91, *p* < 10^−8^) and ECN (t_(61)_ = −4.13, p < 10^−3^) within the listeners’ brains. The aDMN also exhibited negative connectivity with the ECN, but positive connectivity with the LN. The two DMN subsystems showed strong positive functional connectivity (t_(61)_=13.74, *p* < 10^−9^) (Fig. 2B).

#### 3.5 Listener-speaker neural coupling

In all four networks, listeners’ network activities were significantly coupled with the activities of the homologous networks in the speaker’s brain (Fig. 3). Consistent with previous findings (Stephens *et al.* 2010; Liu *et al.* 2017), the coupled activity in the listeners’ brains primarily lagged by that in the speaker’s brain. The temporal profile of inter-brain coupling differed across the four networks. In the LN, the listener’s coupling responses occurred in synchronization with the speaker’s vocalization (0s delay) and peaked with a delay of 4s. In the ECN, aDMN, and pDMN, listener’s coupling responses occurred with a delay of 2s and reached the peak with a delay of 4s, 6s, and 6s, respectively. Among the four networks, the inter-brain coupling in the pDMN was strongest and decayed most slowly. Supplementary analyses demonstrated that the temporal pattern of inter-brain coupling was not accounted for by the autocorrelations of brain network signals (Fig. S4). To validate that the inter-brain coupling in DMN was not an epiphenomenon of task-disengagement, we compared the inter-brain coupling during the first half of communication with that during the second half. We found the coupling strength in the pDMN increased significantly with the unfolding of the communication (Fig. S5).

**Figure 3.**
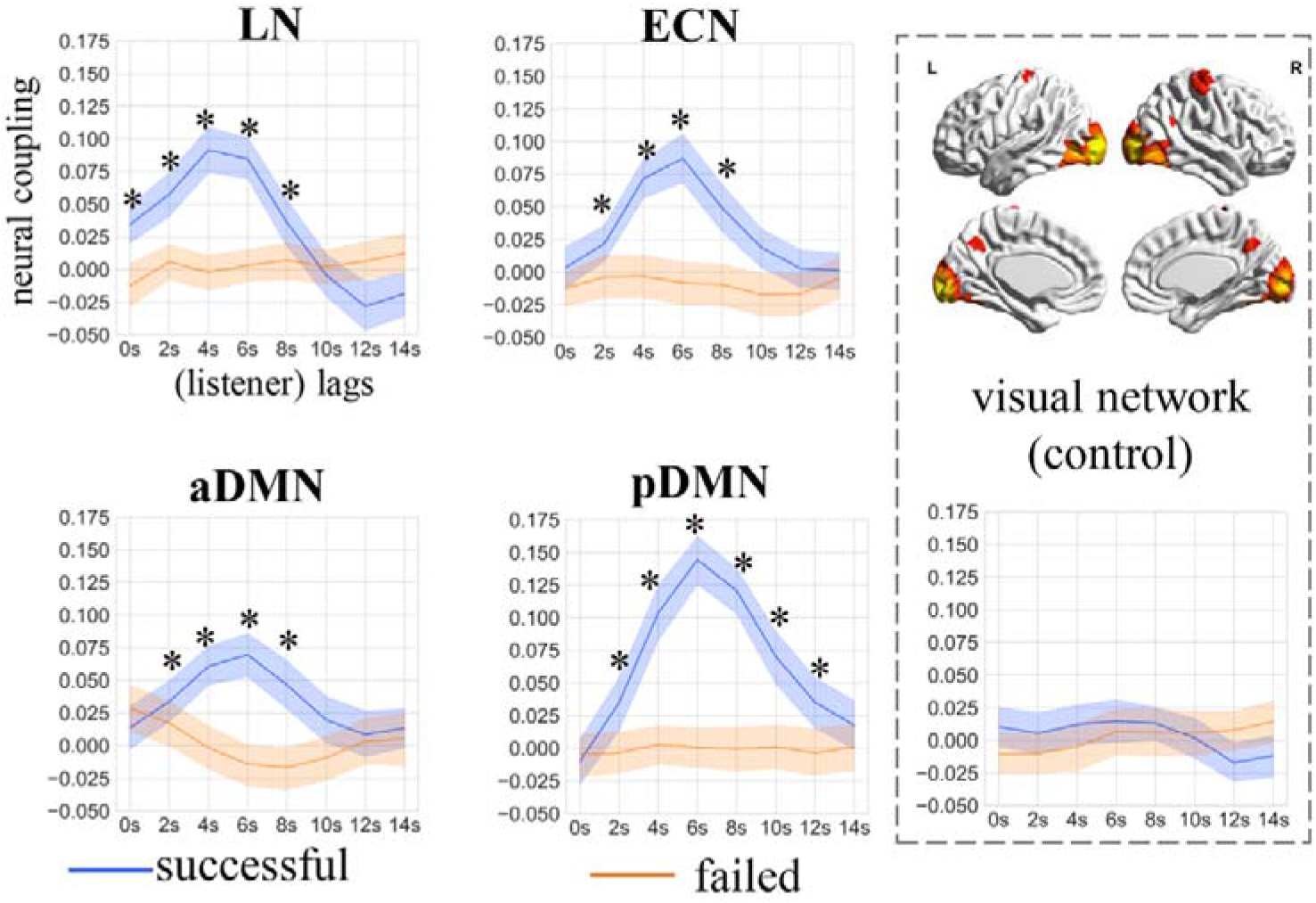
Listener-speaker neural couplings as a function of temporal lags. When communication was successful, listeners’ brain activities in the four networks were significantly coupled with (i.e., correlated over time) the activities of the homologous networks in the speaker’s brain, with several temporal lags. In comparison, no significant inter-brain coupling was found in the visual network. Those inter-brain couplings disappeared when communication failed. Asterisks denote significant differences from zero, with the threshold of *p* < 0.01, FDR corrected.

In comparison, no significant inter-brain coupling was observed in the visual network at any lag, suggesting the inter-brain coupling was specific to task-related brain activities. When communication failed, none of the five networks presented significant listener-speaker neural coupling at any lag, suggesting the observed interbrain couplings were not simply driven by low-level acoustic inputs shared between the speaker and listener. These results also mitigate a methodological concern that the correlation of network time courses between the listener and the speaker was simply resulted from the manipulation that the data from both sides had been pooled together to identify the network component during the group ICA. We note that there was significant ISC among the listeners in the visual network. Moreover, the listeners exhibited significant ISCs in all four networks even when they could not understand the Mongolian story, despite the strength of ISC was significantly lower than that during successful comprehension (Supplementary Fig. S6).

#### 3.6 Listener-speaker neural coupling predicted speech comprehension

To determine the behavioral significance of listener-speaker neural coupling, we examined its association with the listener’s level of speech understanding. A significant positive correlation was found between the coupling strength of pDMN (at lag 6s) and listeners’ comprehension scores (partial r_(60)_ = 0.32, p = 0.012, with memory span regressed out). There was also a positive correlation between the inter-brain coupling in LN (at lag 6s) and comprehension scores (partial r_(60)_ = 0.31, *p*= 0.015, with memory span regressed out) (Fig. 4). No significant correlation was found between the coupling strength of the aDMN or the ECN and the comprehension scores (*p* > 0.1).

**Figure 4.**
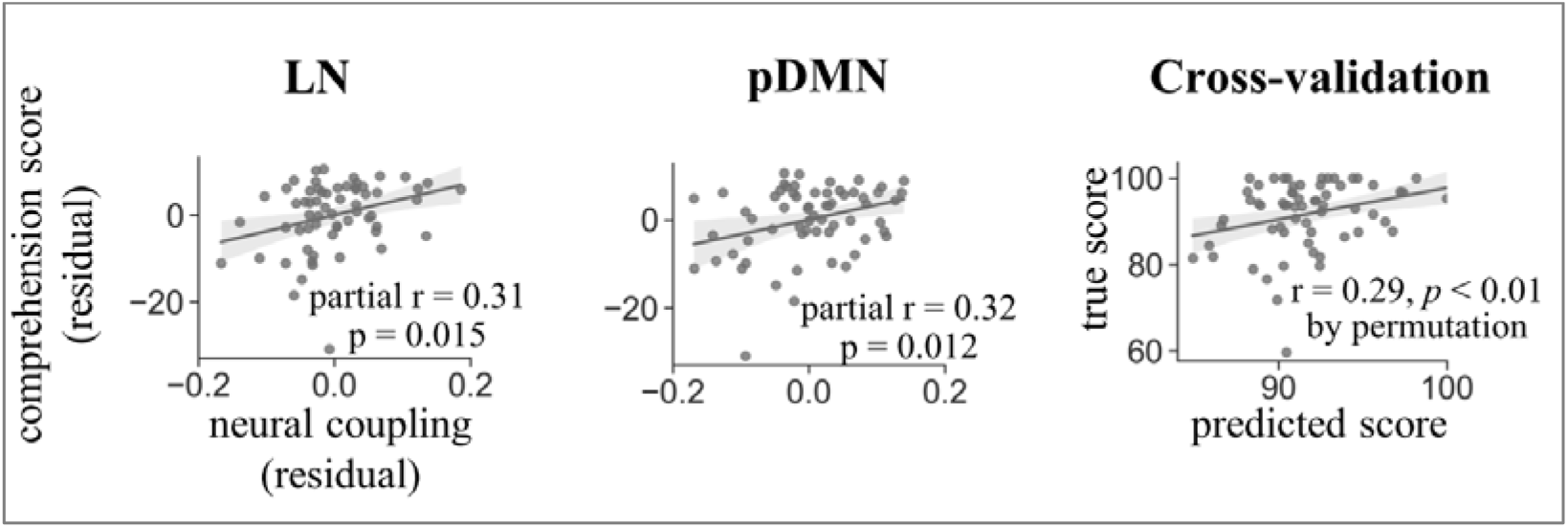
The strength of listener-speaker neural coupling predicts speech comprehension. Left: significant correlations between speech understanding and listener-speaker neural couplings in the pDMN and LN, with memory capacity controlled. Right: Cross-validation for the brain-behavior relationship using a level-one-out procedure.

A leave-one-out cross-validation procedure was performed to evaluate the generalizability of the brain-behavioral relation. In each fold of cross-validation, a regression model was built which took the listener-speaker neural coupling in both the pDMN and LN as inputs and generated predictions of comprehension scores in a novel participant. The predicted scores were positively correlated with actual scores (r_(60)_ = 0.28, p = 0.009) (Fig. 4), with predictive power significantly better than the results from 1000 iterations of permutation tests.

#### 3.7 Inter-network communication revealed by DCM

Results from the two-brain analyses demonstrated that the aDMN, pDMN, LN, and the ECN were actively engaged in language comprehension. We next carried out the DCM analysis to investigate the organization of functional communication among the four networks within the listeners’ brains. The Bayesian Model Selection revealed that model 1 outperformed the other three models. This model hypothesized that the LN and ECN lied in the same layer; both networks then had causal interactions with the aDMN in the intermediate layer, which in turn causally interacted with the pDMN. The posterior probability for model 1 was 0.39, which reflects how likely this model generated the data of a randomly selected subject for this model. The posterior exceedance probability was 0.85, which measured how one model is more likely than any other model (Fig. 5). The group mean of effective connection for the selected model and statistic information were summarized in Table S1.

**Figure 5:**
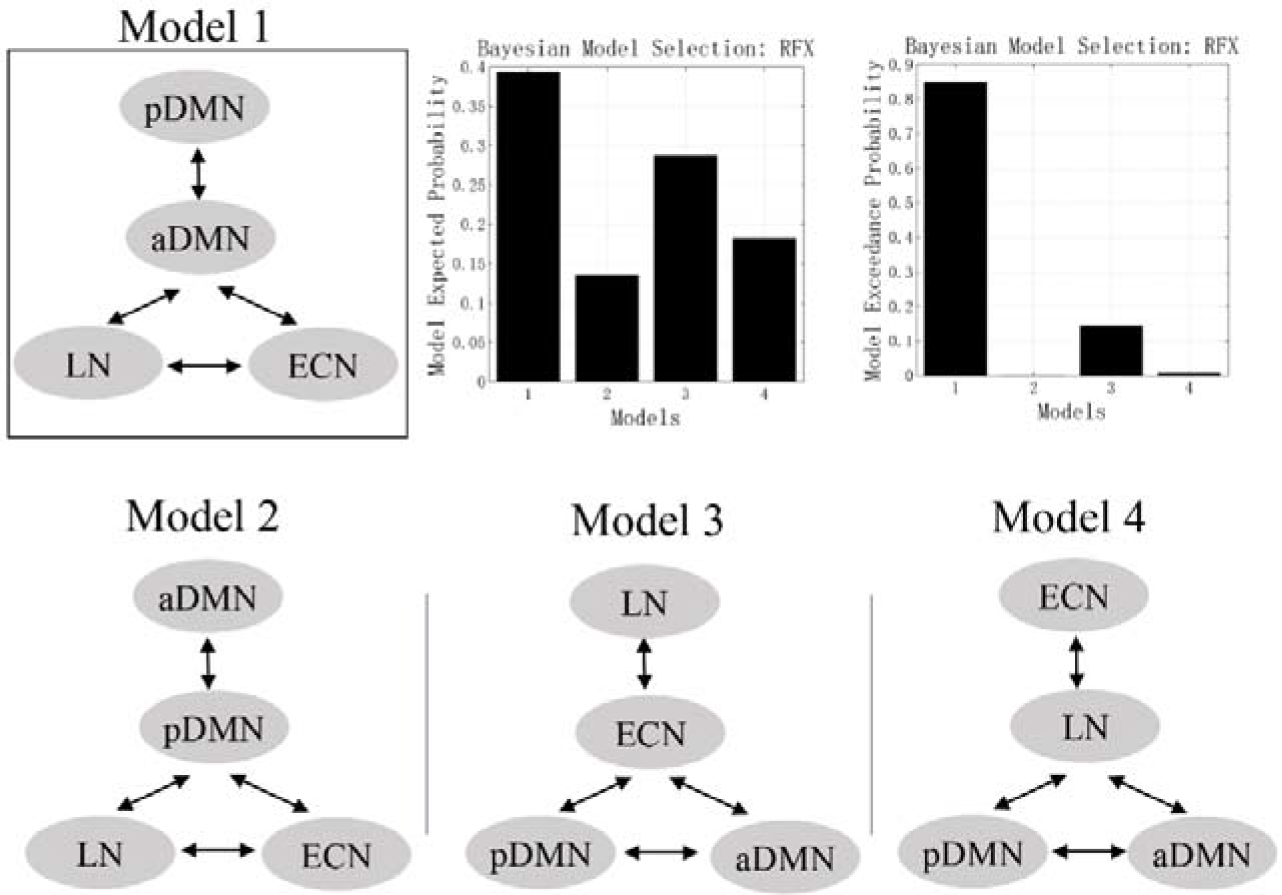
The organization of network communication revealed by Dynamic Causal Modeling. Model 1 was preferred by the random-effect Bayesian model selection.

#### 3.8 Inter-brain coupling in the pDMN increased with greater pDMN-ECN anticorrelation

To gain more insights into the functional properties of DMN, we investigated whether the engagements of aDMN and pDMN in language comprehension (as indexed by the listener-speaker neural coupling) dependents on their interaction with other networks or on the response amplitudes to the speech signals. A significant negative correlation (r_(60)_ = −0.36, p = 0.004) was found between the coupling strength of the pDMN and the pDMN-ECN functional connectivity within listeners’ brains. That is, listeners with greater pDMN-ECN anticorrelation showed stronger neural coupling with the speaker in the pDMN. This association was also found between the change (successful > failed) of pDMN-ECN functional connectivity and the change of pDMN coupling strength (r_(57)_ = −0.32, p = 0.012). There was no significant correlation between the inter-brain coupling and the BOLD signal change in the pDMN (Fig. 6). These results suggest the pDMN’s functionality depends on its interaction with other networks rather than its level of activation. For the aDMN, no significant association between the inter-brain coupling and network connectivity was found, neither between the inter-brain coupling and BOLD signal change.

**Figure 6.**
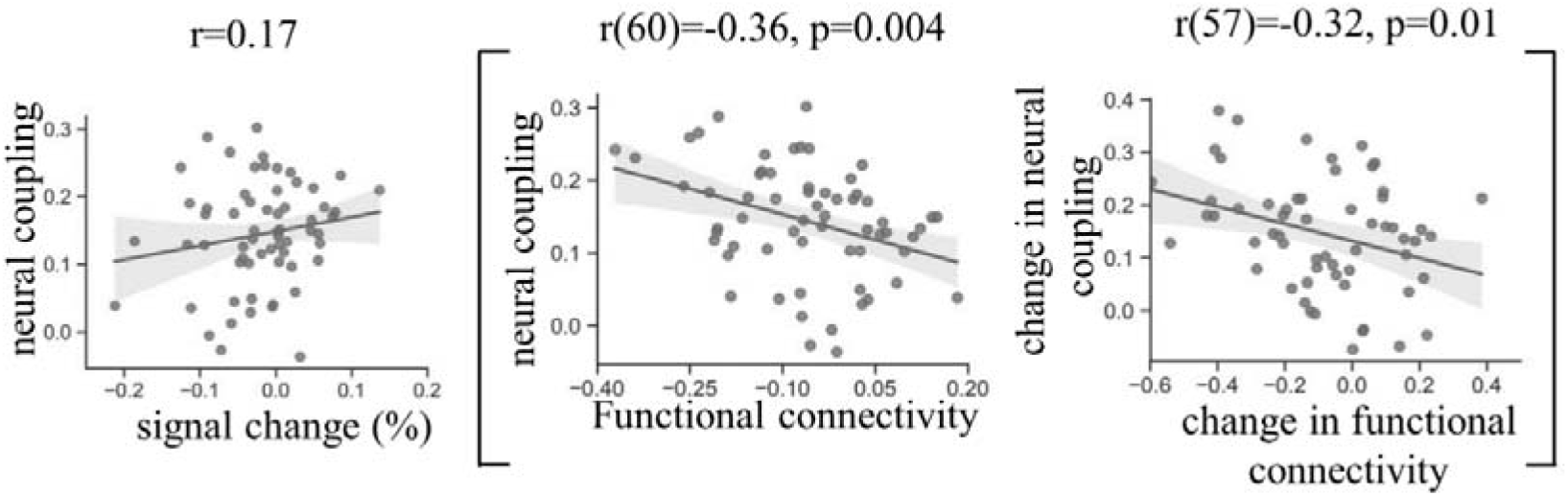
The functionality of pDMN depends on its interactions with other networks rather than its level of activation. Left: Listener-speaker neural coupling in the pDMN was not correlated significantly with the pDMN’s signal change within the listeners’ brains. Right: Stronger listener-speaker neural coupling in the pDMN was associated with greater pDMN-ECN anticorrelation. The change (successful communication > failed) in coupling strength was also correlated with the change in pDMN-ECN functional connectivity.

#### 3.9 Reproducing results with group ICA of different dimensions

To evaluate the robustness of our findings, we reconducted the above analyses using time series derived from a group ICA which separated the spatial map of the brain into 30 independent components. The major results, including the listener-speaker neural coupling, the positive correlation of coupling strength with speech comprehension, the correlation between the interbrain coupling and pDMN-ECN anticorrelation, and the organization of network communication revealed by DCM, are largely consistent across the two cases (Supplementary Fig.S7-S9).

## 4. Discussion

Taking a “two-brain” approach and applying group ICA, this study investigated whether and how the aDMN and pDMN contribute to naturalistic speech comprehension. Consistent with previous findings (Wirth et al. 2011; Rodriguez Moreno *et al.* 2014; Humphreys *et al.* 2015; Jackson et al. 2019), we found the DMN exhibited largely unchanged (in the aDMN) or reduced (in the pDMN) activities while the LN exhibited significantly increased activities during speech comprehension relative to the low-level baseline. Besides, the network dynamics of pDMN were anticorrelated with both the LN and ECN. However, further two-brain analyses revealed that the dynamics of the aDMN and pDMN in the listener’s brain were tightly coupled with the dynamics of the homologous network in the speaker’s brain. Significant listener-speaker neural couplings were also observed in the LN and ECN, but were weaker and decayed faster than that in the pDMN. Moreover, the strength of inter-brain neural coupling in the pDMN and LN predicted the degree to which listeners comprehended the story. In addition, the stronger inter-brain coupling in the pDMN was associated with greater pDMN-ECN anticorrelation. Note, when communication failed, the listener-speaker neural couplings in those networks disappeared; however, there were still significant ISC across the listeners, suggesting the “two-brain” measurement had a higher specificity to task-related neural process. Finally, applying DCM we uncovered a network communication pattern wherein the ECN and LN had direct interactions with the aDMN, which in turn interacted with the pDMN. Implications of these findings are discussed below.

### The active role of “task-deactivated” DMN in speech comprehension

According to the prevalent view, exhibiting activity decrease and anticorrelation with the LN and ECN would indicate that the pDMN interferes with the ongoing speech processing and is suppressed in order to support the externally-directed cognitive processes (Anticevic 2012; Gauffin *et al.* 2013; Zhou *et al.* 2016). In contrast to this notion, the results from the two-brain analyses suggest an active role of the pDMN in spoken narrative comprehension. The inter-brain coupling in the pDMN was unlikely an epiphenomenon of task disengagement, due to the three observations. First, speech production is typically cognitively more demanding than speech comprehension (Hendriks 2014). Therefore, the level of task disengagement in the speaker and that in the listener were unlikely to fluctuate similarly over the course of communication. Even the level of task disengagement covaried to some degree across the speaker and listener, it would most likely result in synchronized rather than listener-delayed coupling in the pDMN as observed in this study. Second, we found the inter-brain coupling in the pDMN during the second half of the communication was significantly stronger than that of the first half. If the pDMN was disengaged in the ongoing task, it should not be influenced by the length of communication. Third, the individual difference in the coupling strength of the pDMN was associated with the individual difference in speech comprehension.

The significant inter-brain coupling and the positive association with task performance demonstrate that the pDMN was not suppressed despite its negative functional connectivity with the ECN and LN. Moreover, we found that the greater pDMN-ECN anticorrelation in the listeners’ brains was associated with stronger listener-speaker neural couplings in the pDMN, indicating that the functional segregation with the executive control system was beneficial for the pDMN’s engagement in speech comprehension. These findings provide empirical evidence supporting the view that, instead of representing simply an antagonistic relationship, network anticorrelations may represent a “division of labor” wherein anticorrelated networks could sometimes work together to complete the same task (Fransson 2006).

Significant inter-brain coupling was also observed in the aDMN. Differing from the pDMN, the aDMN showed no significant change in activity amplitudes during speech listening relative to the fixation period and had positive functional connectivity with the LN. Compared to the pDMN, the inter-brain coupling in the aDMN was weaker, decayed quicker and not significantly correlated with speech comprehension. Consistent with previous studies (e.g., Uddin *et al.* 2009; Seghier and Price 2012b), these results support that the aDMN and pDMN are functionally heterogeneous, possibly engaged in the different stages of language processing with different computations.

### Differences in inter-brain coupling profile among the four networks

In addition to the aDMN and pDMN, we also observed significant listener-speaker coupling in the LN and ECN. In contrast, there was no significant inter-brain coupling in the visual network, suggesting the coupling was specific to task-related neural substrates. Across the four networks, only the LN exhibited inter-brain coupling time-locked to the speaker’s vocalization (i.e., with 0s-lag), probably driven by shared acoustic and perceptual analyses conducted both by the speaker and listener. This is in agreement with the knowledge that (among the four networks) the LN encodes bottom-level perceptual information. Moreover, the coupling strength of the LN was positively correlated with the comprehension score, confirming the critical role of this network in language processing. We also found significant inter-brain coupling in the ECN, supporting the view that successful communication requires the coordination of attention between communicators(Richardson et al. 2007). Together, these results demonstrate that LN, anterior and posterior DMN, and ECN were engaged in narrative comprehension, conforming with the proposed tri-model network of semantic processing (Xu et al. 2017).

Among the four networks, the inter-brain coupling in the pDMN was strongest and decayed most slowly. This distinct feature indicates that the pDMN probably lies at the top of the cortical hierarchy for language processing. Natural communication unfolds over time: the speaker develops a concept and intention, retrieves the lexical-syntactic forms, organizes them into sentences based on grammar principles, and finally produce utterances; the listener analyzes the sounds, maps utterances into meaning, integrates words into sentences, and builds a conceptual-intentional model resembling the speaker’s. Compared to the bottom-level information, the top-level conceptual-intentional information takes relatively longer time for the speaker to encode and for the listener to decode, resulting in the greater delay in the inter-brain coupling of pDMN. Besides, those top-level representations (such as abstract concepts and situation models) may vary less across speech production and comprehension than did the bottom-level representations (such as motor versus acoustic), resulting in stronger listener-speaker neural couplings in the pDMN than in the LN.

### The organization of network communication

The DCM analysis revealed that, compared to the other three possible models, it was more likely that the two externally-oriented networks (the LN and ECN) lied within the same layer, having casual interactions with the aDMN in the intermediate layer, which in turn had direct interactions with the pDMN. This interaction pattern may reflect ordered information flow across the four networks in a hierarchical system. In this system, the LN and ECN occupied the bottom layer, feeding information forward to the aDMN in the intermediate layer, which then feeds information forward to the pDMN in the top layer. Information encoded in the higher layer is also feedback to the lower layer as a “top-down” effect. The suggestion that the pDMN occupied the top layer of the system is in agreement with the inter-brain coupling pattern as discussed in the above paragraph. The suggested hierarchical organization of the four networks is also in line with previous reports on the hierarchical topology of temporal receptive windows (TRW) (Lerner *et al.* 2011). In that study, the early auditory cortices and the mPFC and precuneus were found to be in the low and the top of the TRW hierarchy, respectively. Our assumption is also supported by the observation from a recent TMS study that the nodes in the central executive network can causally regulate activities in the medial prefrontal portion but not the posterior portion of the DMN (Chen et al. 2013). Nevertheless, further studies which directly measure information flow across cortical regions (such as by applying the MEG) are required to test this assumption. In addition, the DCM analysis in this study only tested four possible models of network organization. Future studies should test more possibilities regarding the network organization and include more extensive brain networks in the model.

### The possible mechanism of the DMN’s contribution to language comprehension

Currently, how the DMN may contribute to language comprehension remains elusive. One recent proposal is that the DMN encodes information about the external stimuli and environments that accumulated over a large time scale (Simony et al. 2016). This perspective may still fall in the signal-centered framework. Differing from (but not incompatible with) this view, we speculate that the DMN (mainly the posterior subsystem) may be primarily responsible for conceptual-intentional representations that are internal to organism, as highlighted in the theory regarding the basic design for language faculty proposed by Berwick and colleagues (Berwick *et al.* 2013). In that theory, those internal representations are suggested to be constructed based on the linguistic expressions generated by a basic language module and allow for the semantic-pragmatic interpretation, reasoning, planning, and other activities of the internalized “mental world” (Berwick *et al.* 2013). This mechanism seems to fit well with the observations of the current study. Operating as an internal module, the (posterior) DMN would be highly active during task-free states and not directly modulated by external stimulations, therefore we did not find task-induced activations in this network during speech listening relative to the fixation periods. In addition, computations implemented in the internal module are likely quite different from or even opposite to those implemented in external modules, explaining the anticorrelations of the pDMN with the LN and ECN. Finally, the aDMN in the intermediate layer of the hierarchy may serve as a “connector” that bridges the internal and external systems, thus this network was found to be positively correlated with both the pDMN and LN. We speculate that such an organizational pattern of the four networks may enable the internalization process during successful speech comprehension in which external information is transformed into internal mental representations.

## Conclusions

This study established that, despite showing task-induced deactivation and anticorrelation with the LN the ECN, the pDMN plays an active role in naturalistic speech comprehension, as demonstrated by the significant listener-speaker neural coupling in this network and the positive correlation of network coupling with speech comprehension. Moreover, we demonstrated that the function of DMN mainly depends on its interactions with other networks, rather than its level of activation. Finally, the DCM results together with the inter-brain coupling pattern indicate that the LN and ECN, the aDMN, and the pDMN occupied the bottom, intermediate, and top layers of a hierarchical system, respectively. We conclude that the DMN may primarily work as an internal system that cooperates with the externally-oriented networks, which may allow the transformation of external acoustic signals into internal mental representations during successful speech understanding.

Those novel findings on the functional property of DMN during task processing have important implications for pathological studies. For instance, the overactivation of DMN in elder adults may unnecessarily represent a failure to suppress task-unrelated thoughts, but a shift from a cognitive style dominated by externally-directed processing to a style with more engagement of internally-directed processing. Our study also demonstrates that the “two-brain” approach may provide a unique opportunity for researchers to understand the human brain beyond what can be captured by the “single-brain” approach.

## 5. Acknowledgments

This work was supported by grants from the National Natural Science Foundation of China (NSFC:31900802, 31971036), and the Open Research Fund of the State Key Laboratory of Cognitive Neuroscience and Learning (CNLYB1803). No conflict of interest is declared.

## Reference

Anticevic A. 2012. The role of default network deactivation in cognition and disease. Trends in Cognitive Sciences 16:584–592.

Beckmann CF, Mackay CE, Filippini N, Smith SM. 2009. Group comparison of resting-state FMRI data using multi-subject ICA and dual regression. NeuroImage 47:S148.

Behzadi Y, Restom K, Liau J, Liu TT. 2007. A component based noise correction method (CompCor) for BOLD and perfusion based fMRI. NeuroImage 37:90–101.

Benjamini Y, Hochberg Y. 1995. Controlling the False Discovery Rate: A Practical and Powerful Approach to Multiple Testing. Journal of the Royal Statistical Society: Series B (Methodological) 57:289–300.

Berwick RC, Friederici AD, Chomsky N, Bolhuis JJ. 2013. Evolution, brain, and the nature of language. Trends in Cognitive Sciences 17:89–98.

Calhoun VD, Kiehl KA, Pearlson GD. 2008. Modulation of temporally coherent brain networks estimated using ICA at rest and during cognitive tasks. Hum Brain Mapp 29:828–838.

Carruthers P, Smith PK. 1996. Theories of theories of mind: Cambridge University Press.

Chen AC, Oathes DJ, Chang C, Bradley T, Zhou Z-W, Williams LM, Glover GH, Deisseroth K, Etkin A. 2013. Causal interactions between fronto-parietal central executive and default-mode networks in humans. Proc Natl Acad Sci USA 110:19944–19949.

Davis MH, Johnsrude ISJJoN. 2003. Hierarchical processing in spoken language comprehension. 23:3423–3431.

Dice LR. 1945. Measures of the Amount of Ecologic Association Between Species. Ecology 26:297–302.

Ding N, Melloni L, Zhang H, Tian X, Poeppel D. 2016. Cortical tracking of hierarchical linguistic structures in connected speech. Nat Neurosci 19:158–164.

Fox MD, Snyder AZ, Vincent JL, Corbetta M, Van Essen DC, Raichle ME. 2005. The human brain is intrinsically organized into dynamic, anticorrelated functional networks. Proc Natl Acad Sci U S A 102:9673–9678.

Fransson P. 2006. How default is the default mode of brain function?: Further evidence from intrinsic BOLD signal fluctuations. Neuropsychologia 44:2836–2845.

Friston KJ. 2011. Functional and Effective Connectivity: A Review. Brain Connectivity 1:13–36.

Friston KJ, Kahan J, Biswal B, Razi A. 2014. A DCM for resting state fMRI. NeuroImage 94:396–407.

Friston KJ, Williams S, Howard R, Frackowiak RSJ, Turner R. 1996. Movement-Related effects in fMRI time-series. Magn Reson Med 35:346–355.

Frith CD, Frith U. 1999. Interacting Minds--A Biological Basis. Science 286:1692–1695.

Gauffin H, van Ettinger-Veenstra H, Landtblom A-M, Ulrici D, McAllister A, Karlsson T, Engström M. 2013. Impaired language function in generalized epilepsy: Inadequate suppression of the default mode network. Epilepsy Behav 28:26–35.

Greicius MD, Menon V. 2004. Default-Mode Activity during a Passive Sensory Task: Uncoupled from Deactivation but Impacting Activation. J Cogn Neurosci 16:1484–1492.

Guo CC, Kurth F, Zhou J, Mayer EA, Eickhoff SB, Kramer JH, Seeley WW. 2012. One-year test–retest reliability of intrinsic connectivity network fMRI in older adults. NeuroImage 61:1471–1483.

Hendriks P. 2014. Asymmetries between language production and comprehension: Springer.

Humphreys GF, Hoffman P, Visser M, Binney RJ, Lambon Ralph MA. 2015. Establishing task- and modality-dependent dissociations between the semantic and default mode networks. Proc Natl Acad Sci USA 112:7857.

Jackson RL, Cloutman LL, Lambon Ralph MA. 2019. Exploring distinct default mode and semantic networks using a systematic ICA approach. Cortex 113:279–297.

Kuhlen AK, Allefeld C, Haynes J-D. 2012. Content-specific coordination of listeners’ to speakers’ EEG during communication. Front Hum Neurosci 6:266.

Lerner Y, Honey CJ, Silbert LJ, Hasson U. 2011. Topographic mapping of a hierarchy of temporal receptive windows using a narrated story. 31:2906–2915.

Li W, Mai X, Liu C. 2014. The default mode network and social understanding of others: what do brain connectivity studies tell us. 8.

Li Z, Li J, Hong B, Nolte G, Engel AK, Zhang D. 2021. Speaker-Listener Neural Coupling Reveals an Adaptive Mechanism for Speech Comprehension in a Noisy Environment. Cereb Cortex.

Liu L, Ding X, Li H, Zhou Q, Gao D, Lu C, Ding G. 2021. Reduced listener-speaker neural coupling underlies speech understanding difficulty in older adults. Brain Struct Funct 226:1571–1584.

Liu L, Zhang Y, Zhou Q, Garrett DD, Lu C, Chen A, Qiu J, Ding G. 2020. Auditory–Articulatory Neural Alignment between Listener and Speaker during Verbal Communication. Cereb Cortex 30:942–951.

Liu Y, Piazza EA, Simony E, Shewokis PA, Onaral B, Hasson U, Ayaz H. 2017. Measuring speaker-listener neural coupling with functional near infrared spectroscopy. Sci Rep 7:43293.

McKiernan KA, Kaufman JN, Kucera-Thompson J, Binder JR. 2003. A Parametric Manipulation of Factors Affecting Task-induced Deactivation in Functional Neuroimaging. J Cogn Neurosci 15:394–408.

Menenti L, Garrod S, Pickering M. 2012. Toward a neural basis of interactive alignment in conversation. Front Hum Neurosci 6.

Pamilo S, Malinen S, Hlushchuk Y, Seppä M, Tikka P, Hari R. 2012. Functional Subdivision of Group-ICA Results of fMRI Data Collected during Cinema Viewing. PLoS One 7:e42000.

Richardson DC, Dale R, Kirkham NZJPs. 2007. The art of conversation is coordination. 18:407–413.

Rodriguez Moreno D, Schiff ND, Hirsch J. 2014. Negative Blood Oxygen Level Dependent Signals During Speech Comprehension. Brain Connectivity 5:232–244.

Schoot L, Hagoort P, Segaert K. 2016. What can we learn from a two-brain approach to verbal interaction? Neurosci Biobehav Rev 68:454–459.

Seghier M, Price C. 2012a. Functional Heterogeneity within the Default Network during Semantic Processing and Speech Production. 3.

Seghier ML, Price CJ. 2012b. Functional heterogeneity within the default network during semantic processing and speech production. Front Psychol 3:281.

Shen X, Finn ES, Scheinost D, Rosenberg MD, Chun MM, Papademetris X, Constable RT. 2017. Using connectome-based predictive modeling to predict individual behavior from brain connectivity. Nat Protoc 12:506–518.

Simony E, Honey CJ, Chen J, Lositsky O, Yeshurun Y, Wiesel A, Hasson U. 2016. Dynamic reconfiguration of the default mode network during narrative comprehension. Nat Commun 7:12141.

Smith SM, Miller KL, Moeller S, Xu J, Auerbach EJ, Woolrich MW, Beckmann CF, Jenkinson M, Andersson J, Glasser MF, Van Essen DC, Feinberg DA, Yacoub ES, Ugurbil K. 2012. Temporally-independent functional modes of spontaneous brain activity. Proc Natl Acad Sci USA 109:3131–3136.

Spreng RN, Grady CL. 2009. Patterns of Brain Activity Supporting Autobiographical Memory, Prospection, and Theory of Mind, and Their Relationship to the Default Mode Network. J Cogn Neurosci 22:1112–1123.

Spreng RN, Mar RA, Kim ASN. 2008. The Common Neural Basis of Autobiographical Memory, Prospection, Navigation, Theory of Mind, and the Default Mode: A Quantitative Meta-analysis. J Cogn Neurosci 21:489–510.

Stephens GJ, Silbert LJ, Hasson U. 2010. Speaker–listener neural coupling underlies successful communication. Proceedings of the National Academy of Sciences 107:14425–14430.

Stolk A, Verhagen L, Toni I. 2016. Conceptual Alignment: How Brains Achieve Mutual Understanding. Trends in Cognitive Science 20:180–191.

Uddin LQ, Clare Kelly AM, Biswal BB, Xavier Castellanos F, Milham MP. 2009. Functional connectivity of default mode network components: Correlation, anticorrelation, and causality. Hum Brain Mapp 30:625–637.

Wechsler D. 1987. WMS-R: Wechsler memory scale-revised: Psychological Corporation.

Wilson SM, Molnar-Szakacs I, Iacoboni M. 2007. Beyond Superior Temporal Cortex: Intersubject Correlations in Narrative Speech Comprehension. Cereb Cortex 18:230–242.

Wirth M, Jann K, Dierks T, Federspiel A, Wiest R, Horn H. 2011. Semantic memory involvement in the default mode network: a functional neuroimaging study using independent component analysis. NeuroImage 54:3057–3066.

Xu X, Yuan H, Lei X. 2016. Activation and Connectivity within the Default Mode Network Contribute Independently to Future-Oriented Thought. Sci Rep 6:21001.

Xu Y, He Y, Bi Y. 2017. A Tri-network Model of Human Semantic Processing. Front Psychol 8:1538.

Yan C-G, Wang X-D, Zuo X-N, Zang Y-F. 2016. DPABI: Data Processing & Analysis for (Resting-State) Brain Imaging. Neuroinformatics 14:339–351.

Ye Z, Zhou X. 2009. Executive control in language processing. Neurosci Biobehav Rev 33:1168–1177.

Yu Q, Du Y, Chen J, He H, Sui J, Pearlson G, Calhoun VD. 2017. Comparing brain graphs in which nodes are regions of interest or independent components: A simulation study. J Neurosci Methods 291:61–68.

Zhou L, Pu W, Wang J, Liu H, Wu G, Liu C, Mwansisya TE, Tao H, Chen X, Huang X, Lv D, Xue Z, Shan B, Liu Z. 2016. Inefficient DMN Suppression in Schizophrenia Patients with Impaired Cognitive Function but not Patients with Preserved Cognitive Function. Sci Rep 6:21657.

Zuo X-N, Kelly C, Adelstein JS, Klein DF, Castellanos FX, Milham MP. 2010. Reliable intrinsic connectivity networks: test–retest evaluation using ICA and dual regression approach. NeuroImage 49:2163–2177.

Zwaan RA, Radvansky GA. 1998. Situation models in language comprehension and memory. Psychol Bull 123:162–185.

