## supplementary material for "The “two-brain” approach reveals the active role of task-deactivated default mode network in speech comprehension"

### Assessment for speech comprehension

To make the scoring as objective as possible, we made a list of questions for each story The list included 9-10 questions, which covered important episodes that happened in the different stages of the story (e.g., “please describe the preparation she made for the hiking”, “what happened on her way to the hotel”). Correct answers to these questions contained several key points covering actions, characters, places, time, and motivations et al. For each question on the list, a score was given according to the information a listener provided about those key points. The percentage of the score received out of the total maximal score was computed for each listener.

**Table S1.** The spatial maps of the 16 components from the group ICA not analyzed in the main study.

| IC1 | 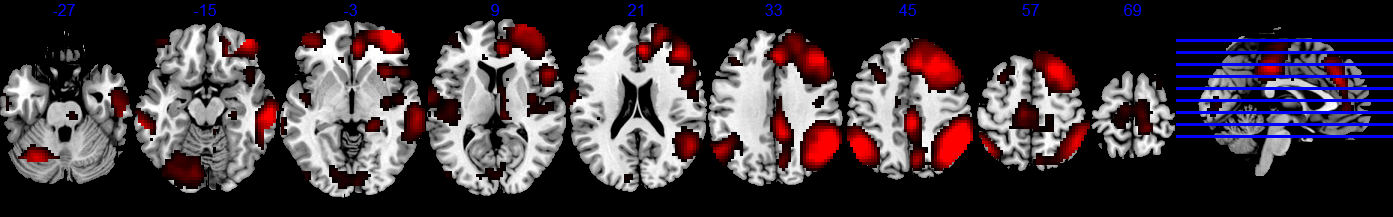 |
| --- | --- |
| IC2 | 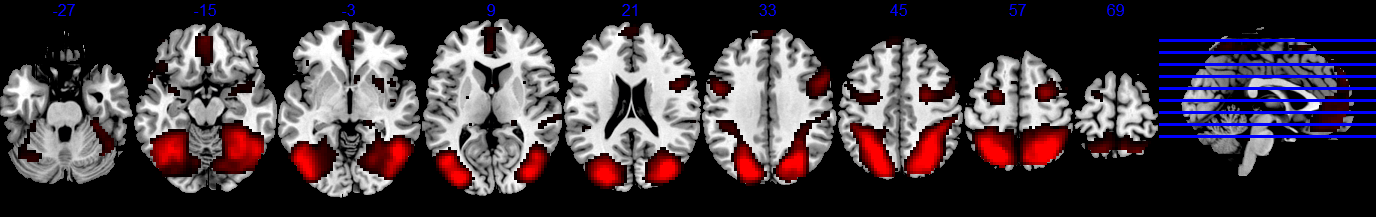 |
| IC3 | 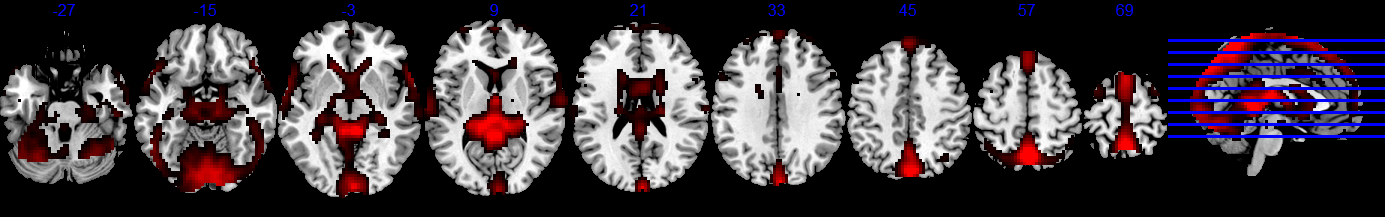 |
| IC4 | 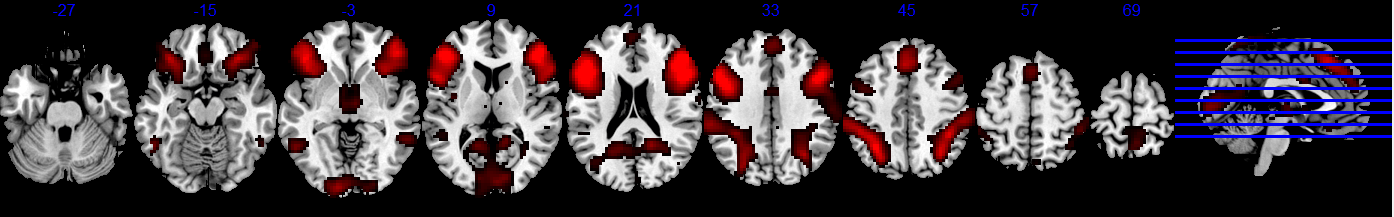 |
| IC5 | 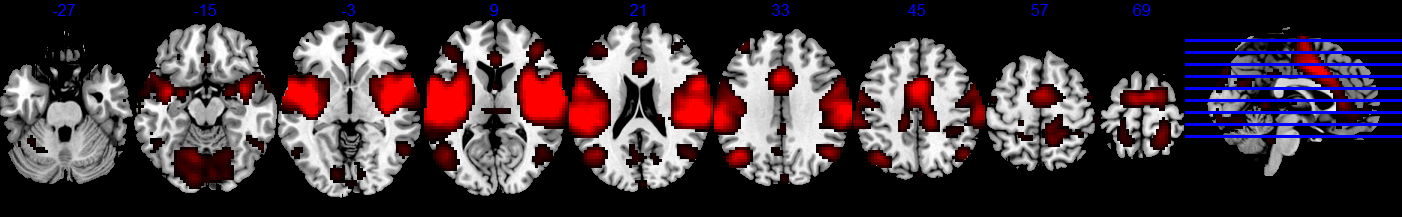 |
| IC6 | 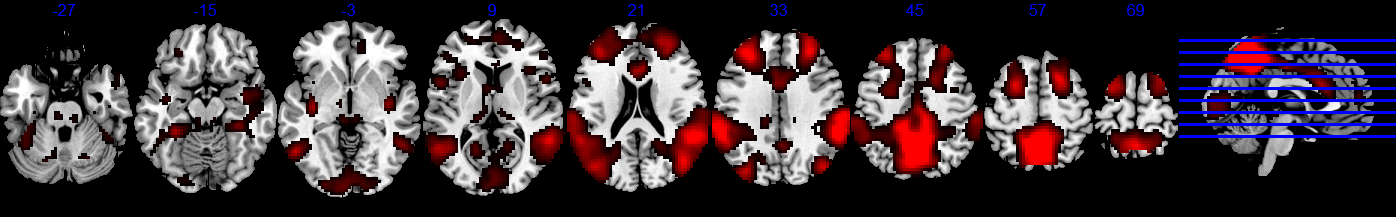 |
| IC 8 | 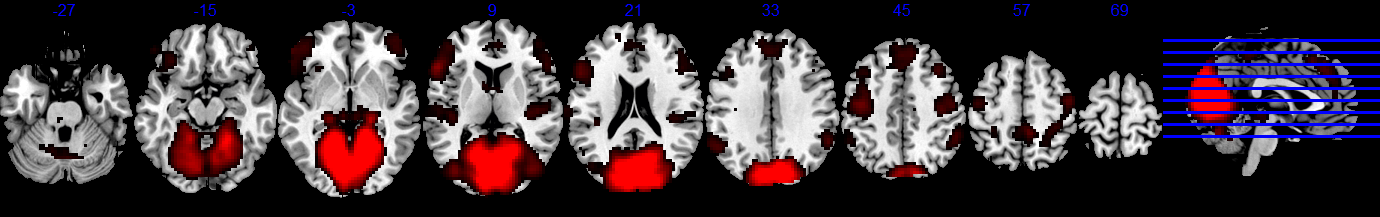 |
| IC 9 | 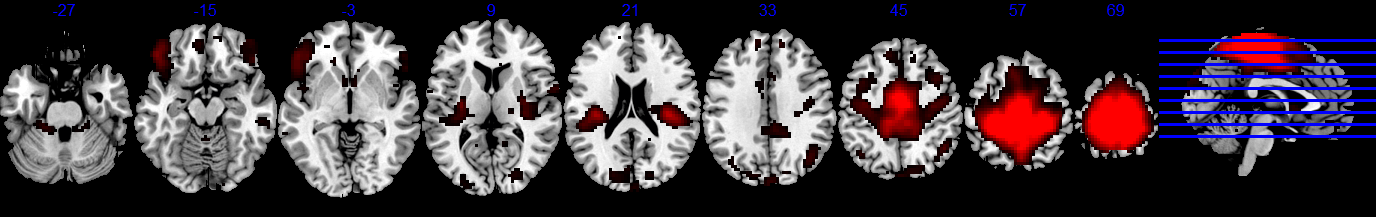 |
| IC10 | 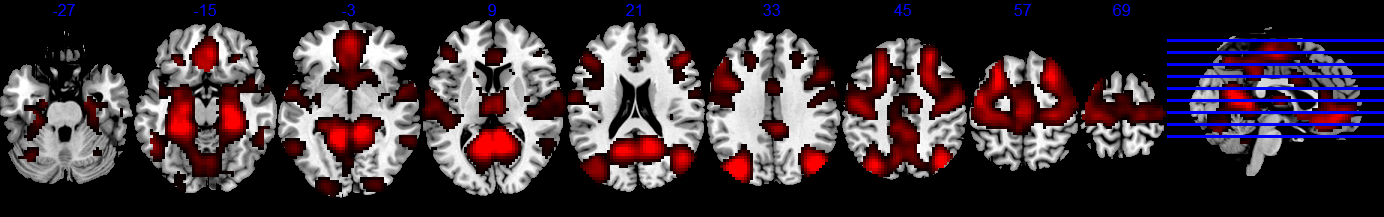 |
| IC12 | 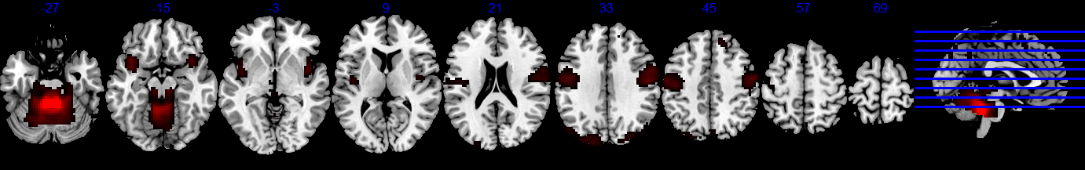 |
| IC13 | 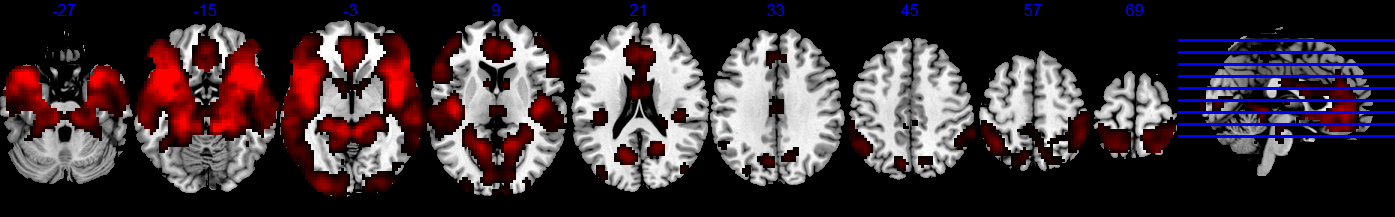 |
| IC14 | 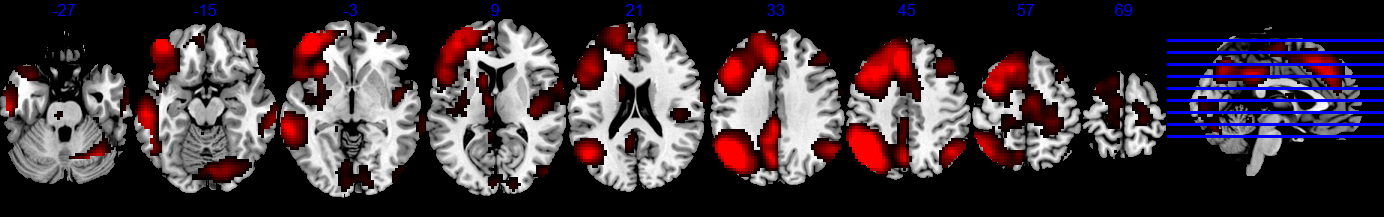 |
| IC15 | 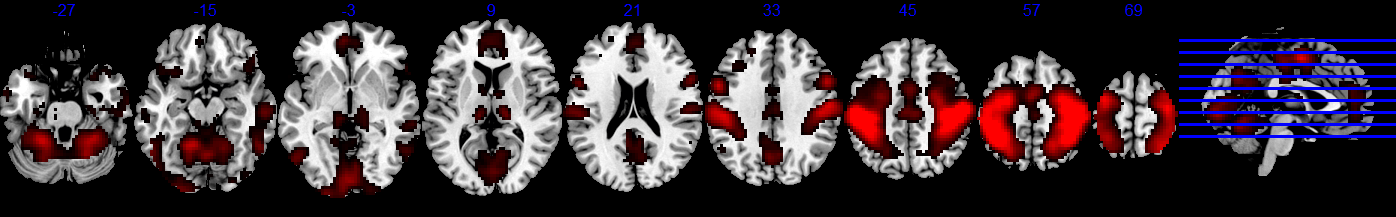 |
| IC17 | 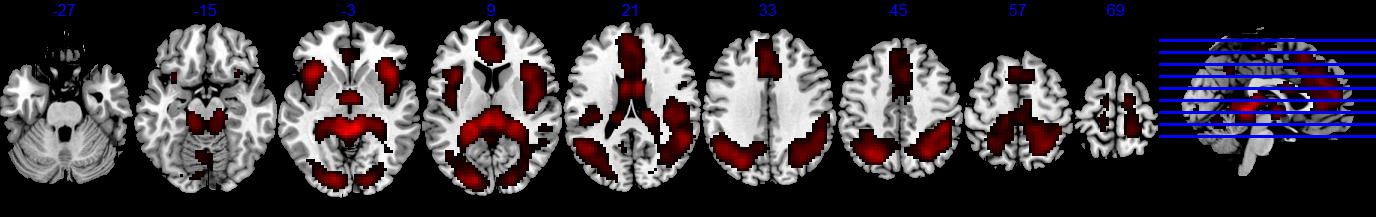 |
| IC19 | 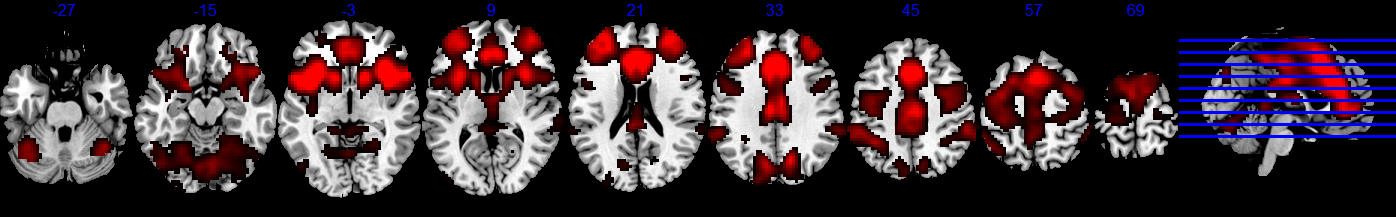 |
| IC20 | 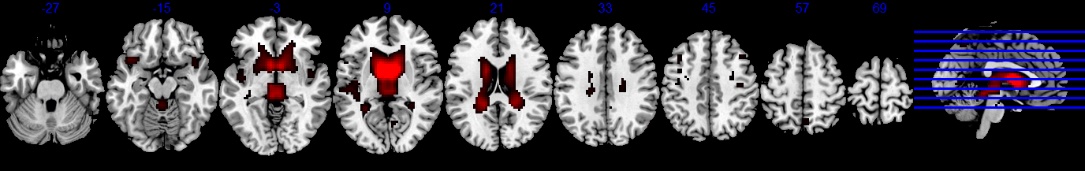  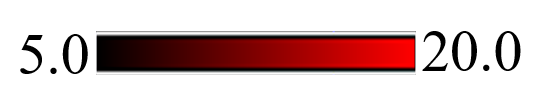 |

### The measurement of spatial overlap using Dice Score

To assess the spatial overlap of component map (i.e., the group t-map) with the meta-analytic map, we first binarized both maps. Then the Dice score was computed using the following equation:

2 * |X ∩ Y| / (|X |+ |Y|)

Where X and Y are the two sets of voxels above threshold in the component map and the meta-analytic map, respectively. |X ∩ Y| means the number of voxels that are common to both sets. |X |+ |Y| means the total number of voxels contained in the two sets.

To reduce bias, we applied a range of thresholds to binarize those maps. Since those meta-analytic maps from the Neurosynth have already been thresholded by an FDR corrected p < 0.05, the threshold range was set to start from the minimum *t*-value in each meta-analytic map to the *t*-value of 6.0, with a step of 0.25. Note, since the overall *t*-values in the IC maps are much higher than the *t*-value in the meta-analytic map, applying the same threshold would produce a large set of above-threshold voxels in the IC map, leading to a relatively low level of spatial overlap between the two types of maps. To alleviate this problem, we adjusted the threshold applied to the IC map. For a given threshold applied to the meta-analytic map (e.g., *t* =3.0), the difference between the averaged *t*-value of above-threshold voxels in the IC map and that of the meta-analytic map was calculated. Then the given threshold plus the global mean difference was used to binarize the IC map.

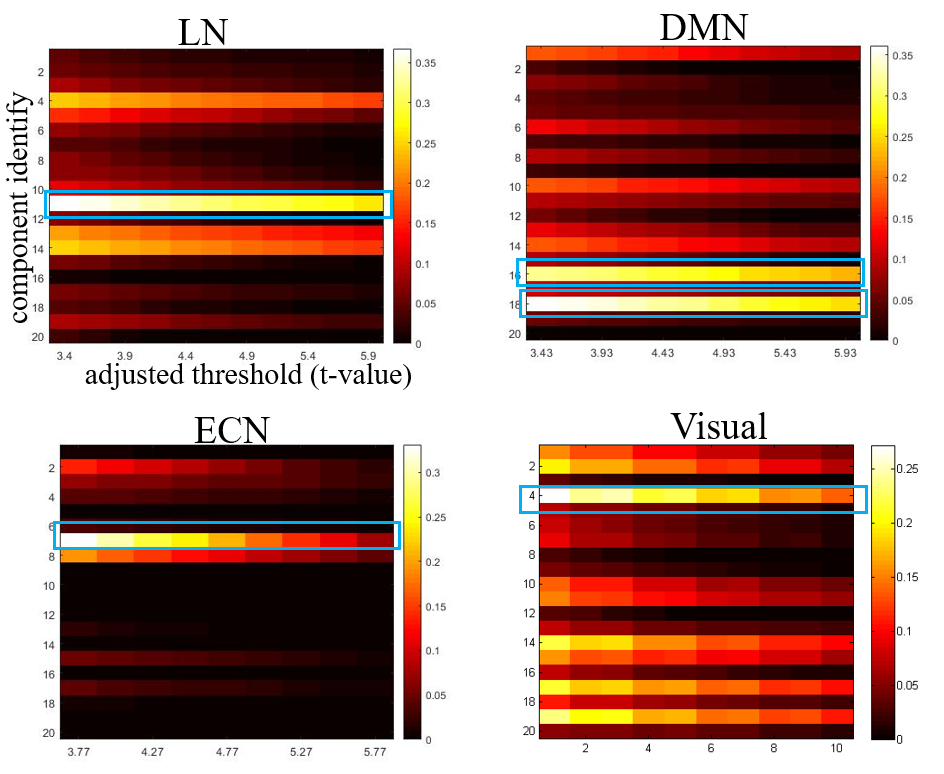

Figure S1: Dice score between each component with the meta-analytic maps. The circled components were chosen to represent the networks of interest.

### Assessing BOLD signal change using ICA-derived time series.

As a supplementary method to assessing BOLD signal changes, we explored ICA-derived time series. For this purpose, we re-conducted the group ICA using the imaging data covering both the 20s resting period and the 10-min story listening period.

Consistent with the distribution pattern of voxel-vise activation, *t*-tests on the signal change calculated using the component time courses revealed significant deactivation in the pDMN (*t*_(61)_ = -2.63, one-tailed *p* = 0.005) and significant activation in the LN (*t*_(61)_= 8.07, one-tailed *p* < 10^-10^) (Fig. 2b). No significant signal change was observed in the aDMN (*t*_(61)_ = -1.04, one-tailed *p* = 0.15) or the ECN (t_(61)_ =1.01, one-tailed *p* = 0.16).

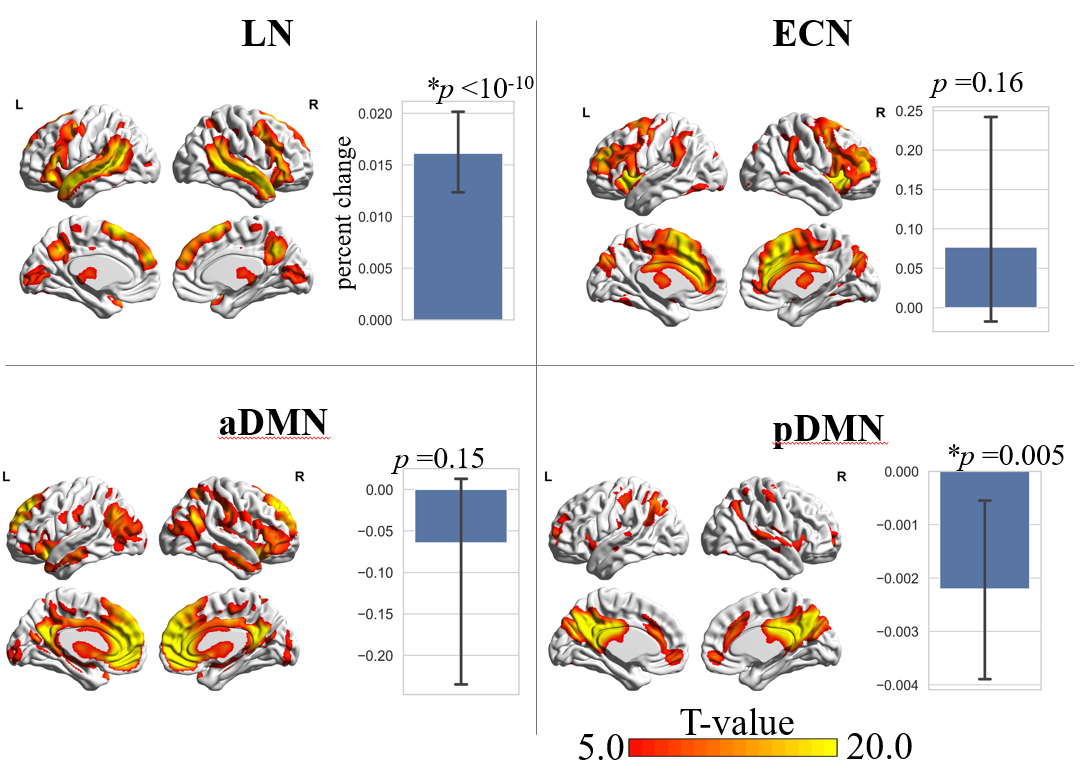

Figure S2. BOLD signal changes in each network assessed using the ICA-derived time courses.

### The effect of baseline length on the activation analysis.

In this fMRI study, 31 participants completed a picture–sound matching task within four task blocks, and viewed scrambled pictures within four baseline blocks. Each block lasted for 30s wherein 15 volumes were acquired. Two methods were employed to examine cortical (de)activations during the task relative to the baseline. In the first method, in line with the analysis reported in the main text, we used only 12 volumes from the first baseline block, and all volumes (N = 60) from the four task blocks to estimate the BOLD signal change. In comparison, a General Linear Model (GLM) was used to assess the (de)activation effect of task > baseline and baseline > task, which included the complete data from all baseline and task blocks. It turns out that, the spatial distributions of activated and deactivated voxels resulted from the two methods were quite similar; while the activation and deactivation effects obtained by the GLM method were statistically more significant.

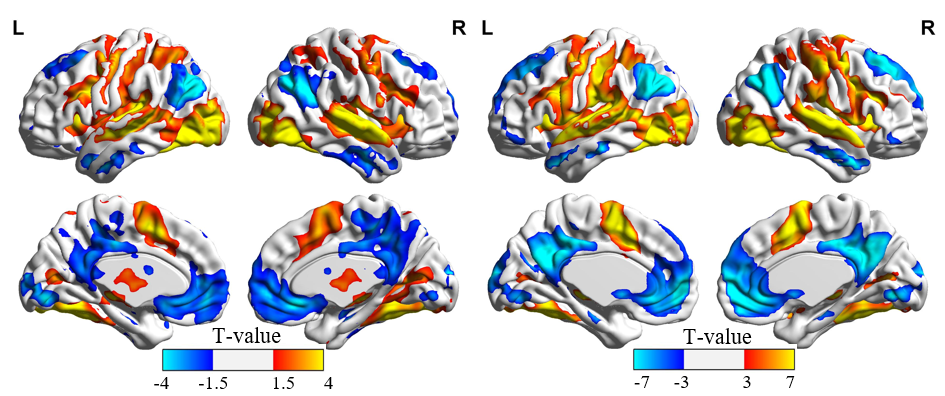

Figure S3. Left: cortical activations and deactivations obtained by the analysis of BOLD signal changes which used only 12 volumes to assess baseline activity. Right: cortical activations and deactivations obtained by the GLM approach which used 60 volumes to assess baseline activity.

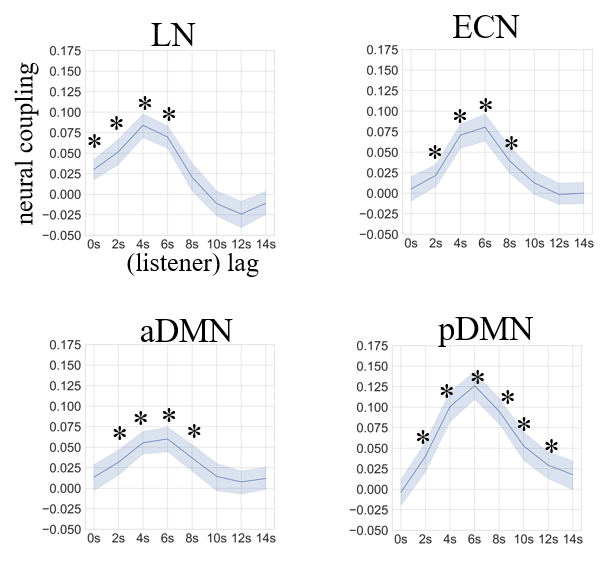

Figure S4. Listener-speaker neural coupling after removing the first-order autocorrelation from the speaker’s time courses.

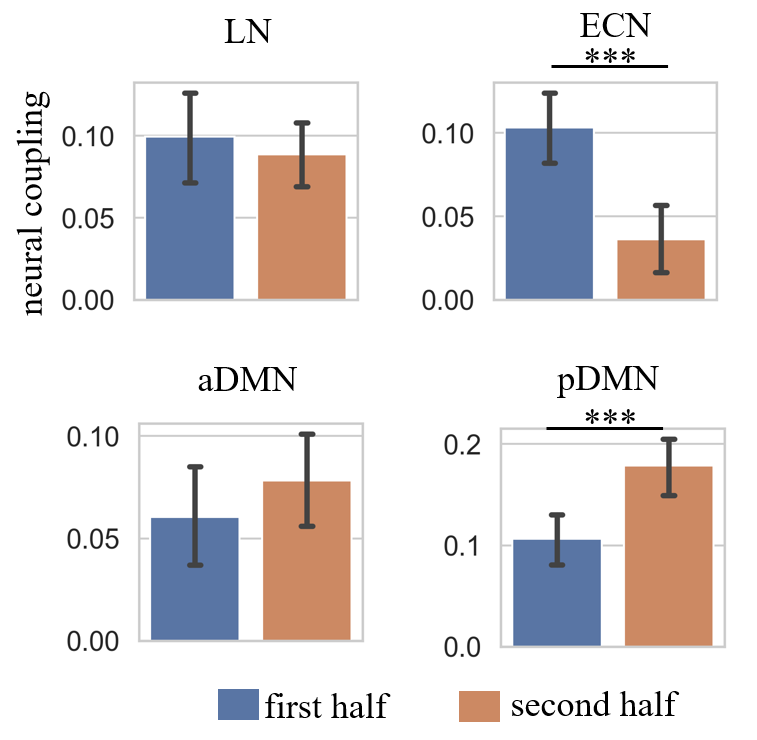

Figure S5. Differences between the inter-brain coupling during the first half (5-min) of communication and that during the second half. Only the peak time for each network was examined.

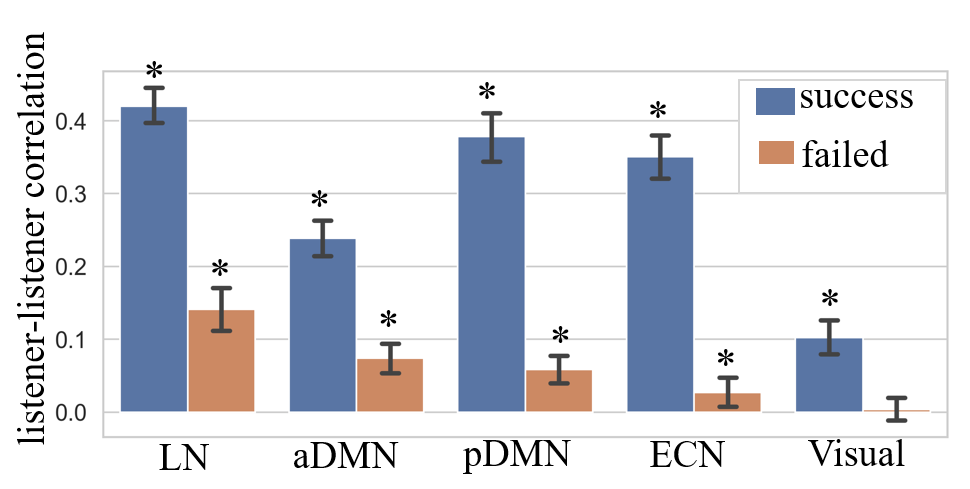

Figure S6. Listener-listener network correlations during the successful and failed communication conditions. For each listener, the inter-subject correlation (ISC) was calculated between his/her network time courses and the averaged time courses of the left 61 listeners. In all five networks, the ISC during the successful condition was significantly higher than that during the failed condition. The asterisk denotes significance at FDR corrected *p* < 0.05.

Table S2. The group mean of effective connections for Model 1 derived from the Dynamic Causal Modelling.

| **To \ From** | **LN** | **pDMN** | **ECN** | **aDMN** |
| --- | --- | --- | --- | --- |
| **LN** | -0.18 *** | 0.00 | 0.04 | 0.05 |
| **pDMN** | --- | -0.22 *** | --- | 0.06 |
| **ECN** | -0.07 ** | --- | -0.04 | -0.21 *** |
| **aDMN** | 0.04 | 0.07 * | -0.03 | -0.21 *** |

*: p<0.05; **: p< 0.005, ***: p < 0.0005, by two-tailed *t*-test.

### Reproducing results with group ICA of 30 dimensions

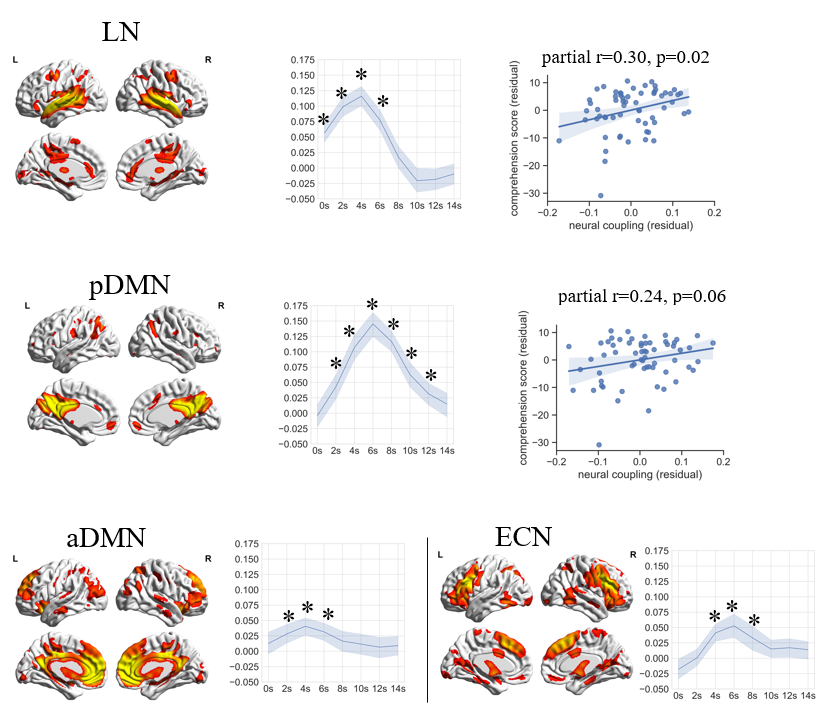

Figure S7. We reconducted a group ICA which separated the whole brain signals into 30 independent components. The spatial distribution of the four networks was quite similar to those derived from the ICA which separating the brain into 20 components. The pattern of lister-speaker neural coupling and the correlation of coupling strength with comprehension score were also consistent across the two cases.

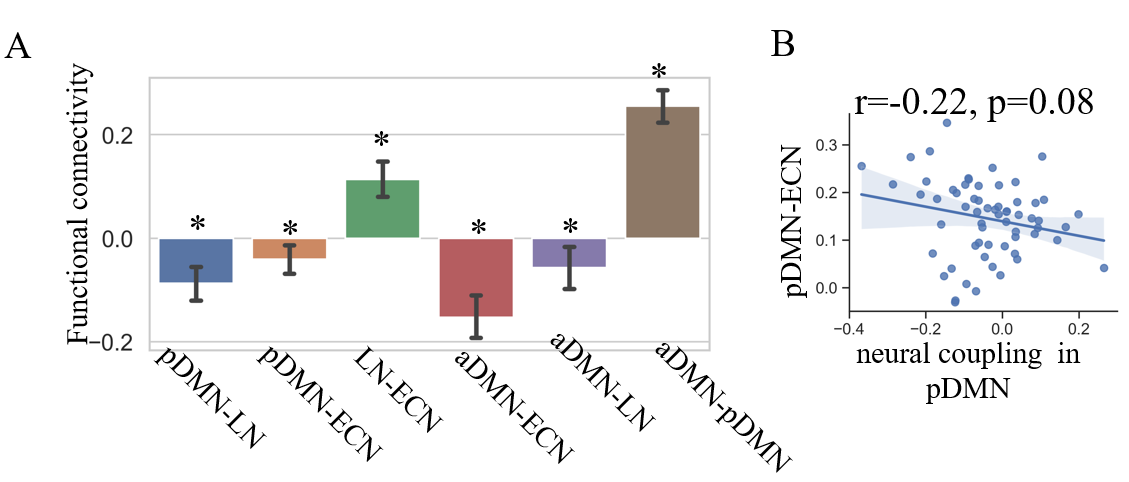

Figure S8. Network connectivity pattern and the association with inter-brain coupling. The time courses used in these analyses were obtained from the group ICA with 30 dimensions.

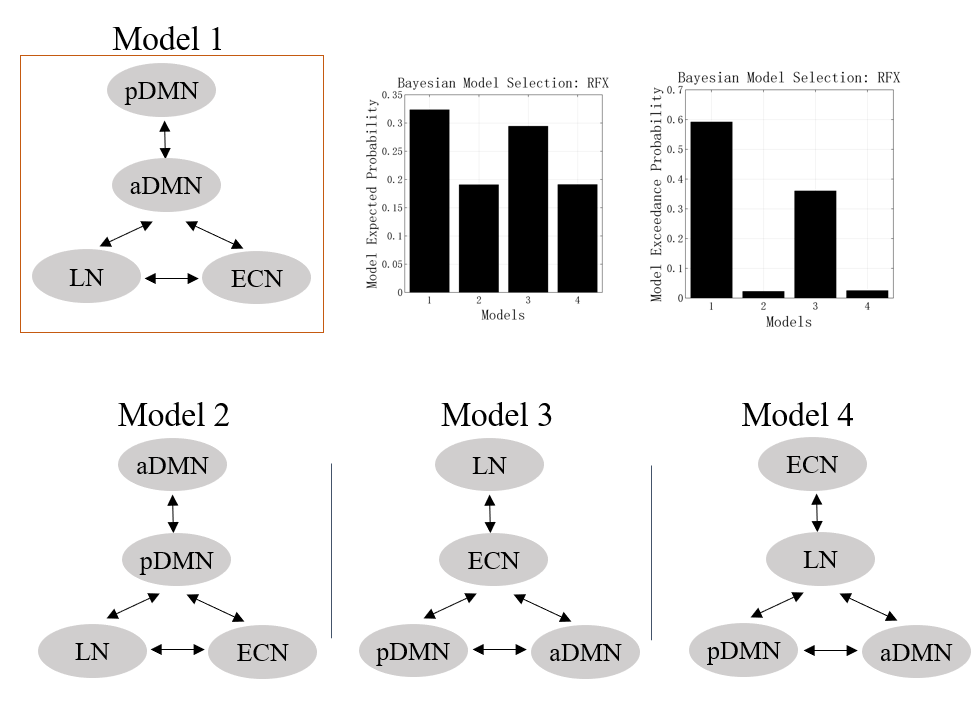

Figure S9. Network organization pattern revealed by Dynamic Causal Modelling. The time courses used in these analyses were obtained from the group ICA with 30 dimensions.
